# Complementary Volume Electron Microscopy-based approaches reveal ultrastructural changes in germline intercellular bridges of *D. melanogaster*

**DOI:** 10.1101/2025.02.18.638836

**Authors:** Irina Kolotuev, Abigayle Williams, Caroline Kizilyaprak, Stephanie Pellegrino, Lindsay Lewellyn

## Abstract

Intercellular bridges are essential to connect developing germline cells. The *Drosophila melanogaster* egg chamber is a powerful model system to study germline intercellular bridges, or ring canals (RCs). RCs connect the developing oocyte to supporting nurse cells, and defects in their stability or growth lead to infertility. Despite their importance, it has been technically difficult to use electron microscopy-based approaches to monitor changes in RC structure during oogenesis. Here, we describe the application of a complementary set of volume EM-based approaches to visualize ultrastructural changes in the germline RCs. The combination of array tomography (AT) and focused ion beam (FIB) scanning electron microscopy (SEM) has allowed us to gain insight into previously unappreciated aspects of RC structure. We were able to quantify differences in RC size and thickness within and between germ cell clusters at different developmental stages. Within a cluster, RC size correlates with lineage; the largest RCs were formed during the first division, and the smallest RCs were formed during the fourth mitotic division. We observed the formation of membrane interdigitations in the vicinity of RCs much earlier than previously reported, and reconstruction of a RC from a mid-stage EC provided insight into the 3D orientation of these extensive cell-cell contacts. Our imaging also revealed a novel membrane structure that appeared to line the interior of the RC lumen. Although the focus was on ultrastructural changes in the germline RCs, our dataset contains valuable details of additional cell types and structures, including the fusome, the germline stem cells and their niche, and the migrating border cells. This imaging framework could be applied to other tissues or samples that face similar technical challenges, where the small structure of interest is located within a large sample volume.

## INTRODUCTION

Intercellular bridges are essential structures that connect somatic cells to each other or developing sperm and eggs to other gametes or to supporting “nurse” cells. Germline intercellular bridges have been observed throughout the animal kingdom from insects to humans; they allow the sharing of haploid gene products, synchronization of meiotic entry, and delivery of materials to support patterning and early embryonic development (Spradling et al., 2022). Although these structures are essential for fertility in many organisms, there is still much to learn about how intercellular bridges are formed, how they are structurally stabilized, and how they grow and expand.

The largest and most well-studied intercellular bridges connect germline cells within the developing egg chamber (EC) of the fruit fly, *Drosophila melanogaster*. The ECs, which ultimately give rise to the mature fly egg, are formed within a structure called the germarium, which lies at the anterior end of the ovariole. Over a dozen ovarioles are found within each ovary; these ovarioles contain a series of ECs at different stages of development, thereby providing a “pseudo-timelapse” of oogenesis within a single field of view (Fig. 1A). EC formation begins with the division of a germline stem cell; one daughter cell, the cystoblast, divides mitotically to generate the sixteen germline cells. These divisions are coordinated across the cluster by a membrane-cytoskeletal structure, the fusome; unequal inheritance of the fusome establishes one of these sixteen germ cells as the oocyte, and the remaining cells will serve as supporting nurse cells (de Cuevas and Spradling, 1998; Deng and Lin, 1997; Diegmiller et al., 2023; Grieder et al., 2000; Lighthouse et al., 2008; Lin et al., 1994; Lin and Spradling, 1995; Snapp et al., 2004). Division of the germline cells is followed by incomplete cytokinesis and stabilization of an intercellular bridge, called a ring canal (RC), which connects the germline cells within the cluster (Ong and Tan, 2010; Robinson et al., 1994; Spradling et al., 2022). Once formed, the RCs must expand nearly 20-fold in diameter, allowing materials to pass from the nurse cells to the oocyte, which is essential to support the production of a viable mature egg.

**Fig. 1.**
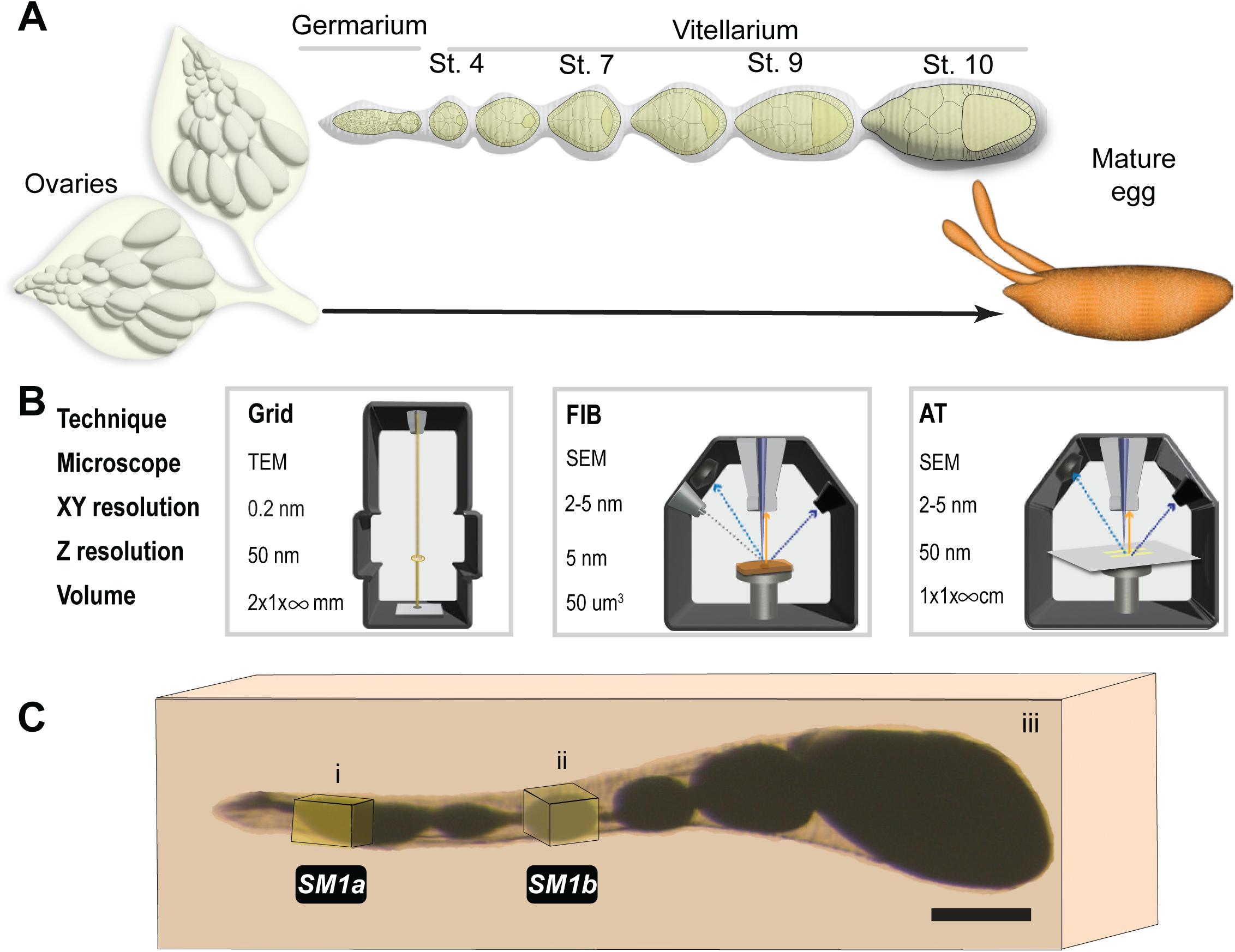
Strategies for sample preparation and analysis of *Drosophila* egg chambers using room temperature electron microscopy. (A) Schematic representation of the organization of the ovarioles within the ovaries. Diagram of a single ovariole showing different stages (germarium through stage 10) of oogenesis before forming a mature egg. (B) Illustration of TEM, FIB-SEM, and AT-SEM approaches. Within each box is an indication of the XY and Z resolution and the potential volume that can be captured by each vEM approach. The TEM microscope uses an electron beam that penetrates through the sample, projecting the image on the camera below. FIB and AT use SEM technology, where only the surface of the sample is scanned by the electron beam. Detectors collect scattered electrons that are “reflected” from the surface after exposure to an electron beam. The difference between FIB and AT is how the external layer of the sample is exposed to the surface. In FIB, the surface is exposed by the gallium ion beam operated inside the SEM sample chamber and subsequently imaged by the scanning electron beam. For AT, embedded samples are sectioned to produce ribbons of sequential sections (arrays) that are transferred to a solid support. Lateral scanning can then be used to identify sections of interest for SEM-based imaging (Fig. S1). (C) Image of an ovariole processed for EM and embedded in epoxy resin. Boxes i and ii, indicate the approximate region of the ovariole that was covered by each of the two FIB-SEM datasets (Supplemental Movies 1a,b) that will be described in subsequent figures. The large box enclosing the entire ovariole (iii) represents the sample covered using AT-SEM. Scale bar is 100 µm. SM refers to corresponding supplemental movies.

The germline RCs are significantly larger and undergo more extensive growth than intercellular bridges that have been studied in other organisms. The requirement for structural stability during regulated growth is likely related to the multi-layered structure of the RCs, which is more complex than the single “rim” that has been reported for other intercellular bridges (Haglund et al., 2011). The germline RCs consist of what is defined as an “inner rim” of actin and actin-binding proteins and a thickened region of the nurse cell membrane, which has been termed the “outer rim”. In addition, extensive membrane invaginations or interdigitations have been observed, at least at the later stages of oogenesis (Loyer et al., 2015; Tilney et al., 1996), but the composition of this region and its role during oogenesis have not been determined. Fluorescence microscopy-based studies have revealed much about RC formation and growth (Gerdes et al., 2020; Hudson et al., 2019; Price et al., 2023; Robinson et al., 1997; Sokol and Cooley, 1999; Tilney et al., 1996; Xue and Cooley, 1993); however, the resolution of light microscopy is insufficient to visualize nanoscale structures. Moreover, because fluorescence microscopy involves the labeling of specific proteins, there are likely other aspects of the developing germline cells that are being overlooked.

Electron microscopy (EM) provides the resolution and cellular context necessary to better understand these essential germline cell structures. Transmission electron microscopy (TEM) is often considered the gold standard of EM, providing Angstrom-level resolution and access to the entire morphome of the cell or tissue. Studies using TEM have revealed aspects of germline cell and RC structure, such as the presence of the membrane interdigitations surrounding the RCs and the changes in actin filament number and length within the inner rim (Loyer et al., 2015; Mahowald, 1971; Riparbelli et al., 2022; Tilney et al., 1996), which would be undetectable by light microscopy. Although a powerful tool, TEM is limited in the sample area and volume that can be imaged (Fig. 1B, S1) and the electron beam penetration. Conventional TEM also suffers from dimensionality reduction, in which three-dimensional (3D) cellular structures are represented as two-dimensional (2D) slices, which can make it challenging to identify or analyze structures of interest without clearly understanding their spatial orientation.

The desire to collect 3D ultrastructural data from tissues and organs has catalyzed a revolution in the EM field, focused on promoting the use and accessibility of volume EM (vEM) approaches (Collinson et al., 2023). vEM includes several powerful methods, many of which use scanning electron microscopy (SEM), to obtain high-resolution images through a large sample volume. Focused Ion Beam (FIB), Scanning Block Face (SBF), and Array Tomography (AT) primarily differ in the way the sample surface is exposed for subsequent imaging, ranging between automatic and manual processing (Figure 1B, S1; Collinson et al., 2023; Peddie et al., 2022). Unfortunately, some experimental questions cannot be completely answered by a single vEM approach; these instead require a carefully selected combination of vEM techniques.

Our desire to understand how the RCs maintain a stable connection between germline cells while also undergoing significant growth motivated us to explore the best EM approaches to monitor RC development during oogenesis. We have considered the strengths and limitations of different vEM approaches, and we have selected a combination of FIB- and AT-SEM to visualize the germline RCs in the developing EC. FIB-SEM provided us with a high resolution, well-aligned dataset that covered multiple young germline clusters, while AT-SEM allowed us to efficiently identify sections that contained germline RCs, from a much larger sample volume, and image these targeted regions at high resolution (Fig. 1B,C). This combination has provided significant insight into the ultrastructural changes that occur during oogenesis. We were able to quantify differences in RC size and thickness within and between germ cell clusters at different developmental stages. Within a cluster, RC size correlates with lineage; the largest RCs were formed during the first division, and the smallest RCs were formed during the fourth mitotic division. We observed the formation of membrane interdigitations in the vicinity of RCs much earlier than previously reported, and reconstruction of a RC from a later stage EC provided insight into the abundance and orientation of these extensive cell-cell contacts. These high-content datasets also revealed a novel membrane structure that appeared to line the interior of the RC lumen. In addition to our findings, this work can serve as an example to guide other researchers who wish to use vEM to study the same or similar tissue types. Although our primary focus was on RC development, these image stacks provide novel insight into the three-dimensional organization of many organelles and structures within the developing EC. To illustrate the power of high-content ultrastructural datasets, we include examples of additional structures in our datasets, the stem cell niche, and the migrating border cells. These datasets can be further mined by researchers studying a variety of topics in oogenesis.

## MATERIALS AND METHODS

### Fly stocks, maintenance, and husbandry

Flies were maintained on cornmeal molasses diet at 25°C. The following lines were used: Cheerio-YFP flies (Kyoto Stock Center #115123), *w^1118^* (Bloomington *Drosophila* stock center, #3605), *matαTub-GAL4* (Bloomington *Drosophila* stock center #7063), and maternal triple driver MTD-GAL4 (*otu-GAL4; nos-GAL4; nos-GAL4;* Bloomington *Drosophila* stock center #31777). The genotype of the flies used for the FIB-SEM dataset was: *esg-GAL4, UAS-hisCFP, Su(H)nlsGFP; ubi-his-RFP/TM6c;* the genotype of the flies used for the AT-SEM genotype: *w^1118^, otu-GAL4; nos-GAL4; nos-GAL4*.

### Immunofluorescence and fluorescence imaging

Prior to dissection, Cheerio-YFP flies (Kyoto Stock Center #115123) or offspring from *w^1118^* crossed to *matαTub-GAL4* were incubated with ground yeast in the presence of males for 44-47 hours at 29°C. Ovaries were fixed in 4% formaldehyde (in PBS) for 15 minutes, washed with PBS + 0.1-0.3% Triton X-100, stained with an Hts antibody (1:20, DSHB 1B1-s) or an HtsRC antibody (1:20, DSHB hts RC-s). Tissue was then stained with an anti-mouse antibody (1:200, Jackson Immunolab), phalloidin (1:500, ECM Biosciences), and DAPI (1:500, D3571 ThermoFisher Scientific). Tissue was mounted in Slowfade Diamond Antifade (Invitrogen) and imaged on a Nikon Ti2-E Inverted microscope equipped with a Yokogawa CSU-X1 Spinning Disk and Hamamatsu ORCA Fusion camera with a 100x Plan Apo VC objective (NA 1.4).

### EM sample preparation

The ECs (FIB-SEM genotype: *esg-GAL4, UAS-hisCFP, Su(H)nlsGFP; ubi-his-RFP/TM6c;* AT-SEM genotype: *w^1118^, otu-GAL4; nos-GAL4; nos-GAL4*) were prepared and fixed using the previously described procedure (Kolotuev, 2014; Loyer et al., 2015). Briefly, the ECs were dissected in PBS and immediately fixed in 2% paraformaldehyde and 2.5% glutaraldehyde in 0.1M PB for two hours at ambient temperature. Samples were postfixed with 1% osmium tetroxide (EMS, 19152) in 1.5% potassium ferrocyanide (EMS-26604). Samples were then incubated in 1% aqueous uranyl acetate solution and dehydrated with increasing concentrations of ethanol (30%, 50%, 70%, 90%, 3 × 100%) and infiltrated with epon-araldite mix (EMS, 1420) resin through increasing percentages of resin with ethanol (Burel et al., 2018; Kolotuev, 2014).

### Block preparation and ultramicrotomy

Samples were flat embedded in epon-araldite and polymerized at 60°C for 48 hours, following the procedure described previously (Kolotuev, 2014). The flat embedding is crucial for the later orientation of the blocks for sectioning (Fig. 1C, S1A). After polymerization, samples were trimmed to remove empty resin and to expose the area of interest with a 90° diamond trim tool (Diatome AG, Nidau, Switzerland) and mounted on a Leica UC6 microtome for trimming (Leica, Austria).

#### FIB-SEM Sample Preparation

Samples were trimmed and attached to aluminum stubs using a silver epoxy mix (Kizilyaprak et al., 2019) while carefully orienting the surface of interest facing the edge (Fig. S1). The samples were oriented perpendicular to the sectioning plane to ensure the smallest dimension was created for imaging (∼60 µm wide, Fig. S1).

#### AT-SEM Sample Preparation

Samples prepared for AT-SEM were sectioned using the ultramicrotome as described (Burel et al., 2018; Franke and Kolotuev, 2021; Kolotuev, 2014). Briefly, polymerized flat blocks were secured inside the ultramicrotome holder and carefully oriented perpendicular to the sectioning plane with the overall ROI slightly protruding from the tip (Fig. S1). After trimming, a dedicated ats knife (Diatome AG, Nidau, Switzerland) was used to generate the sequences of consecutive sections, or arrays. Typically, 50-200 sections were collected per ribbon with 50-150 nm thickness. Arrays were transferred to 2×4 cm bits of a silicon wafer (Ted Pella, No 16015), and wafers were air-dried and placed in an oven at 60°C for one hour for additional adhesion (Fig. S1).

### EM acquisitions

For both types of acquisitions, the HELIOS Nanolab scanning microscope (Thermo Fischer Scientific) was used. Images were acquired with a backscattered electron detector (MD) at an accelerating voltage of 2 kV, probe current 0.8 nA at a 2.5mm distance from the detector. To achieve a TEM-like appearance, the micrographs were collected in the inverted mode.

#### FIB-SEM

The acquisitions were performed automatically and piloted by the Slice and View program (Thermo Fischer Scientific), as previously described (Kizilyaprak et al., 2019). The 10.2 nm-pixel resolution stack (germarium) was acquired at 25 nm cutting thickness (the raw data pixel size of 0.0102 × 0.0102 × 0.025 µm^3^), and the 9.9 nm-pixel resolution stack (stage 4 EC) was acquired at 20 nm cutting thickness (the raw data pixel size of 0.0099 × 0.0099 × 0.020 µm^3^).

#### AT-SEM

The acquisitions were performed using the Maps program (Thermo Fischer Scientific) in semi-automatic acquisition mode as single frames or stitched mosaic panels to cover more extensive regions like those previously described (Franke and Kolotuev, 2021). After lateral screening for specific features of interest, mid-range resolution (∼20 nm-pixel size) images were acquired to identify the ROI (Burel et al., 2018) more precisely. Then, the ROIs were collected with higher resolution (5nm-pixel size) as single images or tiles of consecutive images. Acquisition parameters were adjusted based on the dataset, varying between low resolution/high acquisition speed parameters during screening and high resolution/low acquisition speed during detailed maps and stack acquisition.

### Alignment, Segmentation, and Image Analysis

To align and segment the datasets, a combination of different programs was used including Fiji TrakEM2 (Cardona et al., 2012), IMOD (Kremer et al., 1996), Amira (Thermo Fischer Scientific), and Adobe Photoshop. The specific approach was selected based on the quality of the stack and the required alignment precision. FIB-SEM datasets do not require much alignment, as the image stacks are obtained, with small z-steps and little distortion. With AT-SEM, there is some distortion due to the sectioning limitation, thicker z-sections, and overall larger acquisition area/volume. In this case, 3D alignment is more challenging and, in some cases, multiple programs were required to obtain a well-aligned stack.

Manual segmentation of the samples was done using the IMOD 3dmod program; the contours were traced by hand and extrapolated to generate the 3D model. The recognition of the structures of interest was done by the operator. The structures that could not be traced manually due to their complexity were rendered using the isosurface function of 3D mod (https://bio3d.colorado.edu/imod/doc/3dmodHelp/modvIsosurface.html). This tool uses a selected sub-volume to generate surfaces in the model from regions where the intensity value crosses a threshold.

## RESULTS

### Analysis of the germline ring canals requires volumetric, high-resolution imaging

EC formation begins at the anterior end of the germarium, where a germline stem cell divides to produce a daughter cystoblast. This cystoblast then undergoes four synchronized mitotic divisions, producing a cluster of 16 germline cells. These divisions are not complete, and a stable intercellular bridge, or RC will connect the daughter cells. Once formed, the compact germline cell clusters (Fig. 2A) advance in their development as they move away from the stem cell niche. Although TEM studies have provided insight into RC structure and function, a systematic analysis of ultrastructural changes in the RC during development has not been performed. The spatial distribution and random orientation of these small structures (0.5-10 µm in diameter) mean that single 50-100 nm sections will not capture the entire ring. Moreover, the “end-on” view through the central profile of this toroid structure is extremely rare (Fig. 2B). Instead, the appearance of the size and thickness of the same RC can differ depending on the orientation of the section (Fig. 2C, Supplemental Movie 2). Therefore, only by acquiring, superimposing, and even segmenting the entire sample volume, can the total structure be analyzed reliably (Fig 2B).

**Fig. 2.**
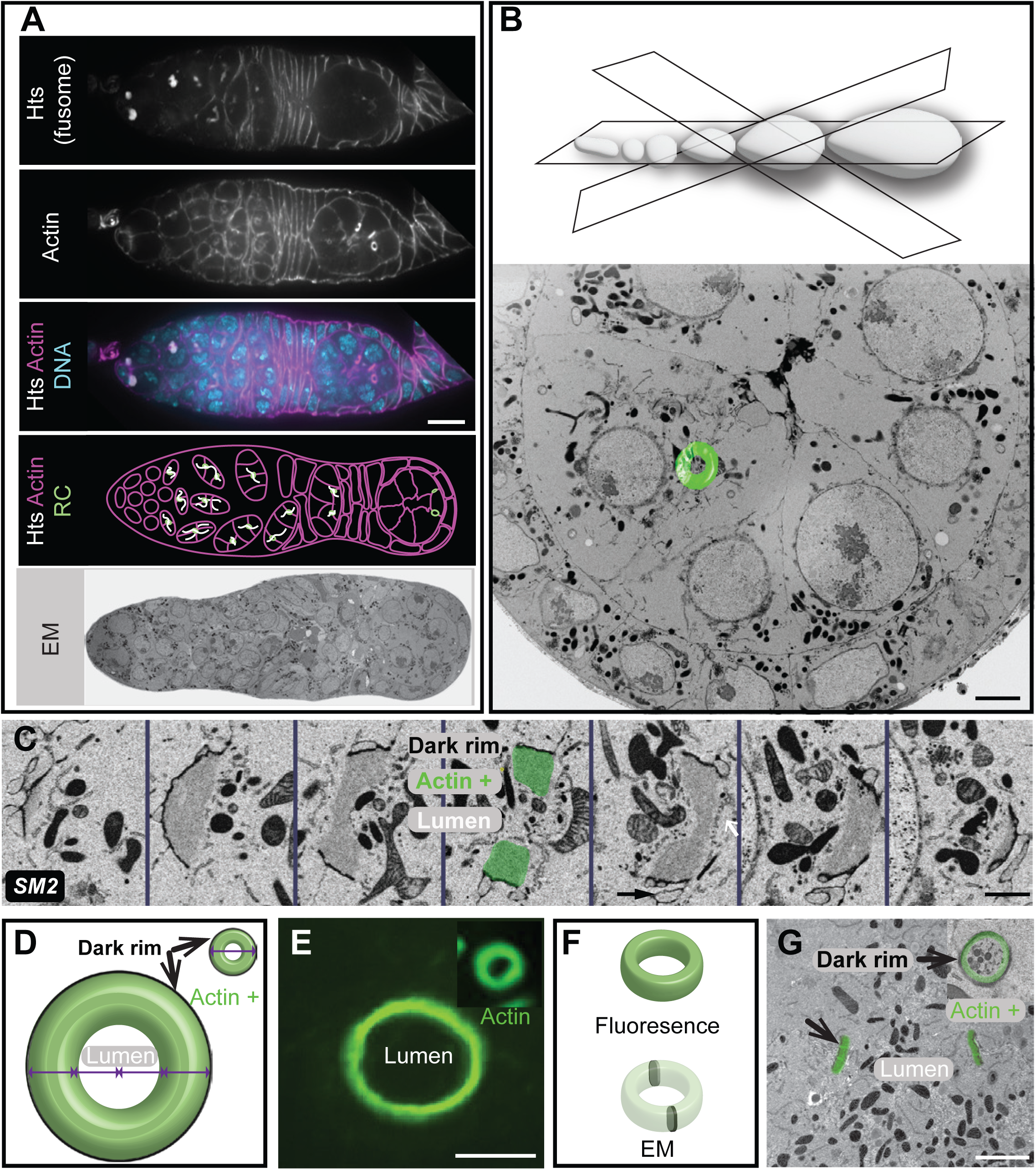
*Drosophila* egg chamber development and germline ring canals. (A) Single-section confocal fluorescence images of the germarium stained with an Hts antibody to label the fusome (white), phalloidin (magenta), and DAPI (cyan). Scale bar 10 µm. An image of a single section through the germarium of the AT-SEM image shows the richness of an EM dataset but also highlights the complexity of the structure. (B) Different sectioning orientations result in the highly variable appearance of the same structure, which is why vEM (shown in green overlay) provides the most informative view of these structures. Scale bar 3 µm. (C) FIB-SEM images of different sections through the same RC (from a stage 4 EC) demonstrate the different ways that the same RC can appear. Scale bar 1 µm. (D) Cartoon illustrates how the size of the RC changes dramatically in size and volume during oogenesis. (E) Maximum intensity projection of 3 sections taken of a RC from a stage 1 (inset) or stage 10 EC stained with an HtsRC antibody taken at the same magnification. Scalebar is 5 µm. (F) Diagram showing the structure of the RC and how it might be observed by fluorescence microscopy or a single EM section. (G) AT-SEM images of RCs taken from germarium (inset) or a stage 8 EC. Scale bar is 1 µm. SM refers to corresponding supplemental movie.

As oogenesis proceeds, the overall size of the EC increases, making it ever more difficult to locate and image the entire RC within older ECs (Fig. 2D-G). For example, a random thin section that might have been sufficient to partially capture several RCs at the early stages of oogenesis will likely not contain any at later developmental time points. Therefore, the visualization of a single RC might require between 20 and 200 consecutive sections, depending on the stage of the EC and sectioning orientation. For our analysis, we decided on a combination of two vEM approaches, FIB-SEM and AT-SEM, to visualize RC development throughout oogenesis.

### FIB-SEM reveals early ultrastructural differences in ring canal size and thickness within germline cell clusters

Because the germline cells divide in a stereotypical pattern, the lineage of each germline cell and connecting RC can be assigned. For example, a single RC is formed after the first mitotic division (M1), two are formed after the second mitosis (M2), four are formed after the third mitosis (M3), and eight are produced by the final mitotic division (M4; Fig. 3A). Only a few studies have described lineage-based differences in ring canal size or structure. One TEM-based study reported that within the germarium, RC diameter and accumulation of material in the inner rim correlated with lineage (Koch and King, 1969); however, a detailed analysis of this ultrastructural variation was not performed. A more recent light microscopy-based study reported lineage-based differences in the degree of contractile ring constriction; with each germline cell division, the contractile ring constricted further, such that the ring canals that formed from the first mitotic division had a larger starting diameter compared to those formed following the subsequent divisions (Ong and Tan, 2010). Recently, our group has demonstrated that there are lineage-based differences in RC scaling. Specifically, the largest M1 RCs increase in size more slowly than the smaller RCs derived from later mitotic divisions (Shaikh et al., 2024). Our high-resolution FIB-SEM datasets provided us with the opportunity to further explore the lineage-based ultrastructural differences and develop testable hypotheses to explain the observed differences in RC scaling.

**Fig. 3.**
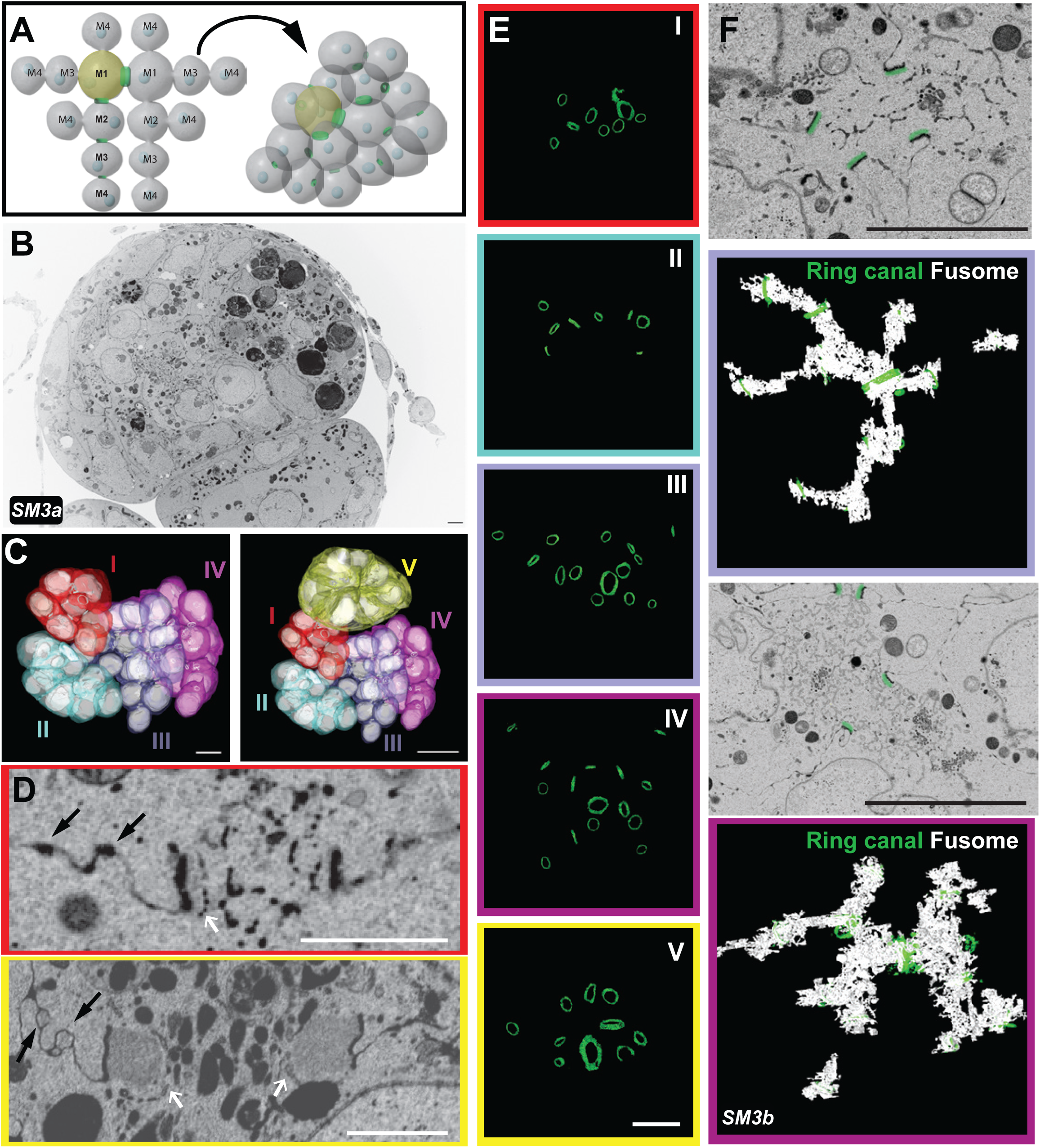
FIB-SEM analysis of multiple germline cell clusters within the germarium reveals differences in ring canal size and structure. FIB-SEM was used to acquire 10.2 nm resolution images of part of the germarium. (A) Diagram shows the stereotypical organization of the germ cell cluster as it often appears in drawings (left) compared to the way that it is packaged within the tissue (right), which highlights the importance of rendering. (B) Single section from the FIB-SEM stack shows the content-rich images. (C) Rendering of the germline cell membranes and nuclei within all clusters contained in the stack. Scalebars are 5 µm (left) and 10 µm (right). (D) Single sections of RCs from one of the youngest (red) and the oldest (yellow) germline cell clusters demonstrate the differences in RC diameter and thickness. White arrows point to the membrane structure lining the ring canal lumen; black arrows point to electron-dense regions of the nurse cell membrane and membrane invaginations. Scale bars are 1µm. (E) The isosurface tool was used to render the RCs from all clusters shown in (C). These 3D models allow us to visualize the differences in RC size, thickness, and spatial orientation within each cluster. Scale bar is 5 µm. (F) Single sections from the FIB-SEM image stack showing cross sections of the fusome passing through multiple RCs (pseudo-colored in green). Rendering of the fusome using the isosurface tool generates a model of the branched fusome structure in two germ clusters (III and IV) from region 2b of the germarium. Scalebar is 5 µm. SM refers to corresponding supplemental movies.

Due to the compact nature of the germarium, we expected that a single FIB-SEM dataset could contain many germline cell clusters (Fig. 1C, box i, Supplemental Movie 1a), allowing us to compare RC size and toroid thickness within and between different germline cell clusters. The first FIB-SEM dataset was acquired at ∼10 nm XY- and 25nm Z-resolution (Fig. 3B, Supplemental Movie S3a). To establish the developmental sequence and to understand the tissue “landscape,” we used the IMOD program to manually segment the contours of germline cell membranes and the nuclei within those cells (Kremer et al., 1996). We could identify five germline cell clusters at different developmental stages; we color-coded them based on their connections, which allowed us to determine the lineage and position of cells within each cluster (Fig. 3C, Supplemental Movie S3a). Although we do not have complete volumes for all clusters, we were able to conclude that two clusters (I and II) are located within region 2a of the germarium, two clusters are within region 2b (III and IV), and one cluster is in region 3/stage 1 (Fig. 3C). We found that the RCs within the younger germ cell clusters (I and II) were smaller in diameter and appeared thinner than those in the older germ cell clusters (III-V). The average RC diameter measured from the darker outer rim was 1.27 +/- 0.28 µm in region 2a (clusters I and II), 1.35 +/- 0.32 µm in region 2b (clusters III and IV), and 1.79 +/- 0.61 µm in region 3/stage 1 (cluster V). Even though our measurements are precise, they are not absolute since the tissue tends to shrink ∼10% during the dehydration and embedding steps. These measurements are larger than some previously reported based on measurements from fluorescence images, but the lineage and stage-based differences in the ratio are consistent with prior studies (Mahowald and Strassheim, 1970; Ong and Tan, 2010; Shaikh et al., 2024).

Single sections through the center of the RC allowed us to visualize the ultrastructural details of each RC within the germ cell clusters. From these images, we could easily define the thickness of the actin-rich rims and the structure of the darker outer rims. Outside of the outer rim, we observed additional electron-dense regions along the cell membrane, reflecting areas of stronger cell-cell adhesion (Fig. 3D). Near these electron-dense regions, we often observed membrane invaginations, which became more obvious within the older germ cell clusters (Fig. 3D; black arrows point to regions of stronger cell-cell adhesion and membrane invaginations). These regions of the membrane were reminiscent of the extensive invaginations, or “microvilli,” which have been observed by single-section TEM of later-stage egg chambers (Loyer et al., 2015; Tilney et al., 1996). To our knowledge, this is the first time that these membrane structures have been observed in younger germline cell clusters.

To segment the structurally complex RCs, we used the IMOD isosurface tool. This nonselective tracking method often results in a less precise tracing of the structure of interest, tracking all structures of a similar threshold within the analyzed area. Therefore, in addition to the toroid components, the material passing through the rings and in their periphery was included in the rendering data. Despite that, we could visualize the position, relative size, and orientation of all RCs within the data set (Fig. 3E; Supplemental Movie S3a).

While analyzing RCs in different clusters of the germarium, we could not ignore the presence of the fusome. The fusome is an essential and conserved germ cell structure composed of membrane and cytoskeletal elements, such as Hts (Fig. 2A). Of the sixteen germline cells, two contain four RCs; of those two pro-oocytes, the cell that inherits a larger portion of the fusome will become the oocyte, suggesting that the fusome also plays a role in oocyte determination (de Cuevas and Spradling, 1998; Deng and Lin, 1997; Grieder et al., 2000; Lighthouse et al., 2008; Lin et al., 1994; Lin and Spradling, 1995; Spradling et al., 1997). Despite its importance, very little analysis of the fusome has been done with vEM. Because this structure was obvious in our FIB-SEM dataset, we used many individual isosurface volumes to render the entire fusome from the most well-sampled clusters in the dataset, cluster III and cluster IV (both from region 2b of the germarium). In Region 2b, the mitotic divisions have ended, and the fusome is breaking down. In cluster III, the fusome still passes through all visible RCs. In cluster IV, which may be slightly older, we can still see it passing through or near all but one of the fifteen RCs. Both clusters show regions of discontinuity within the germline cells that are not directly connected to the pro-oocytes (Fig. 3F, Supplemental Movie S3b). Additional analysis of this dataset or new FIB-SEM datasets could be performed to fully characterize the assembly and breakdown of this essential germline cell structure.

In contrast to fluorescence microscopy, which is used to highlight specific proteins or structures of interest, EM reveals the surrounding cellular context without specific labeling, which can provide unexpected insight into structures of interest. For example, our analysis detected ER-like membrane structures lining many RCs, starting from the stages containing the fusome, but still visible after its breakdown (Fig. 3D, white arrows). Although EM only provides a single snapshot of the tissue, these membrane structures did not appear to be passing through the RCs; instead, they seem to cap the inner rim of actin (Fig. 3D, S4C). Future studies can further explore the identity of this membrane structure and whether it might play a role in stabilizing the germline RCs.

### FIB-SEM revealed lineage-specific differences in ring canal size and thickness

In addition to the differences in RC size between clusters of different developmental stages, our analysis revealed obvious differences in the size and thickness of RCs within each cluster. It has previously been shown that at least beginning at stage 3, there is a hierarchy to nurse cell size; the four nurse cells directly connected to the oocyte are larger than the cells that are separated from the oocyte by two RCs, and so on (Diegmiller et al., 2021; Imran Alsous et al., 2017). One possibility is that RC size also varies based on distance from the oocyte; if this were the case, then the RCs that connect the first layer of nurse cells directly to the oocyte could be the largest, and the RCs would become progressively smaller as they are further away from the oocyte (Diegmiller et al., 2021; Imran Alsous et al., 2017). Alternatively, RC size could be based on lineage. It has been demonstrated that the RCs that are formed from different mitotic divisions constrict to different endpoints, with the RC that originated during the first mitotic division (M1) reaching the largest final diameter and the RCs that originate from the second, third, and fourth mitotic divisions (M2, M3, and M4 respectively) reaching progressively smaller final diameters (Ong and Tan, 2010). Therefore, we could use vEM to compare RC size and thickness based on location and lineage within the germline cell clusters.

To gain further insight into the inter- and intra-cluster differences in RC size and thickness, we acquired a FIB-SEM dataset of an older EC (stage 4; Figure 1C, box ii, Supplemental Movie 1b) obtained from a deeper level of the same sample. The EC at this stage is larger than the germarium; due to the technical limitations of the FIB-SEM microscope, we were unable to capture the entire EC volume. We were, however, able to image portions of all nurse cells, the entire oocyte, and eleven RCs (Supplemental Movie 4). With the two vEM datasets, we could use the slicer tool to orient all RCs in the same way and compare RC structure. We found that lineage was a more reliable predictor of RC size and thickness. In cluster III, the average RC size ranged from 1.07 +/- 0.14 µm for the M4 RCs to 2.43 µm for the M1 RC. A similar trend was observed in cluster IV (Fig. 4, S4; 1.21 +/- 0.09 µm for M4 RCs to 2.03 µm for the M1). For cluster V, we did not have full coverage of the entire EC volume, so we could not definitively determine the lineage of each RC (Fig. S4); however, for those we could reliably categorize, we found that the M4 RCs ranged from 1.31-1.40 µm, and the M1 RC was 3.26 µm in diameter (Fig. 4, S4). The size of the lighter electron density actin within the inner rim also varied based on lineage. In clusters III and IV, the M1 RC contained a thin layer of actin just interior to the electron-dense outer rim; at this stage, the other RCs (M2-M4) did not show any apparent actin accumulation. By stage 1, an inner rim of actin was obvious in all RCs, but the thickness of that ring varied significantly based on lineage (Fig. 4, S4). These data demonstrate the importance of matching not only the developmental stage but also the RC type when comparing experimental phenotypes, as differences in RC size or thickness could be due to inherent differences in their structure and not due to specific perturbation or genetic background.

**Fig. 4:**
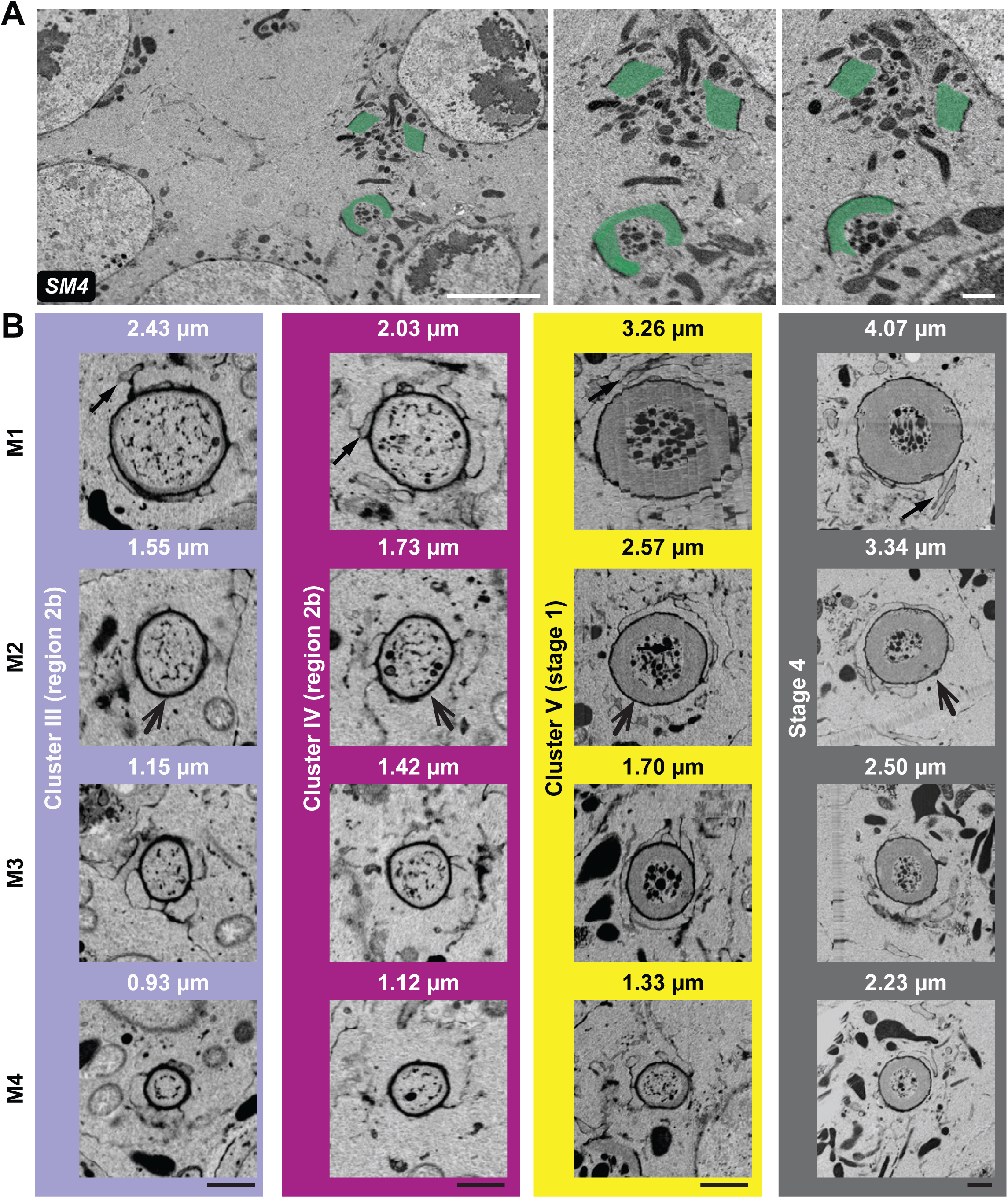
Ring canal ultrastructural differences are best appreciated in 3D. (A) Single FIB-SEM sections through the same two RCs from a stage 4 EC. The actin-containing inner rim is pseudocolored green. The size and thickness of the ring can appear different depending on the orientation of the RC within the section. Scalebars are 5 µm (left) and 1 µm (right). (B) The Slicer tool was used to generate end-on views of RCs from each cluster; M1-M4 indicates the mitotic division of origin of each RC. Scalebars are 1 µm. SM refers to corresponding supplemental movie.

Despite the obvious differences in the absolute thickness of the inner rim, if we compare thickness to RC diameter, the differences are not as dramatic, at least for the M1-M3 RCs. For example, in the stage 4 EC, the thickness of the inner rim was 24-29% of the RC diameter for the M1, M2, and M3 RCs but was only 19% for the M4 RC. This pattern was also observed in the stage 1 EC, where the thickness of the inner rim was ∼21-23% of the diameter of the rings in the M1, M2, and M3 RC, but it was thinner and more heterogenous in the M4 RC (ranging from 4-12% depending on where the measurement was made). This suggests that the differences in RC size and structure observed in the germarium (Fig. 3, S4; (Ong and Tan, 2010) are maintained at least through stage 4 of oogenesis. Despite the differences in the size and thickness of the inner rim, we did not observe detectable differences in the thickness of electron-dense region of the ring, or the outer rim. The resolution of the SEM data (5-10 nm) did not allow us to measure the nuanced differences in the thickness of the outer rim, so additional analysis of this part of the structure will require a more sensitive approach, such as TEM.

### Array Tomography-SEM provides a straightforward approach to locating and imaging ring canals and membrane interdigitations within older egg chambers

FIB-SEM provided insight into ultrastructural differences in the RCs (∼1-4 µm in diameter) within developing germline cell clusters and young egg chambers, which occupy a relatively small sample volume (∼50 µm^3^). Ultrastructural analysis of later developmental stages, where the RCs are distributed within a much larger volume, cannot be efficiently performed using this approach. AT-SEM provides a more straightforward approach to easily locate the non-uniformly oriented RCs within a much larger sample volume while maintaining the ability to collect vEM data (Fig. 1B). Using this approach, we collected longitudinal sections through an ovariole (the entire germarium through stage 10) onto five wafers; each wafer contained 200-300 100 nm-thick sections (Fig. 5A-C). The ribbon on most wafers remained intact, making it possible to define the precise axial sequence of the sections. Although the ribbon of sections on one wafer was disrupted (Fig. 5B; wafer 4), all sections were collected, so the sequence could be re-established during imaging or in post-acquisition processing. The strength of the AT approach lies in the ability to collect data in stages; low-resolution maps of each wafer identify the relevant sections and areas of interest for subsequent higher-resolution imaging (Fig. 5D,E, Supplemental Material 5). Because FIB-SEM allowed us to capture ring canals within the germarium and in early oogenesis, our primary goal was to visualize RC structure in the middle and later-staged ECs (Fig. 5E, Supplemental Movies 5a,b). Following longitudinal screening of the wafers, we located sections containing RCs and acquired either higher magnification tiled images (Fig. 5F) or a sequence of sections for 3D analysis (Fig. 5G,H; Supplemental Movies 5a,b).

**Fig. 5:**
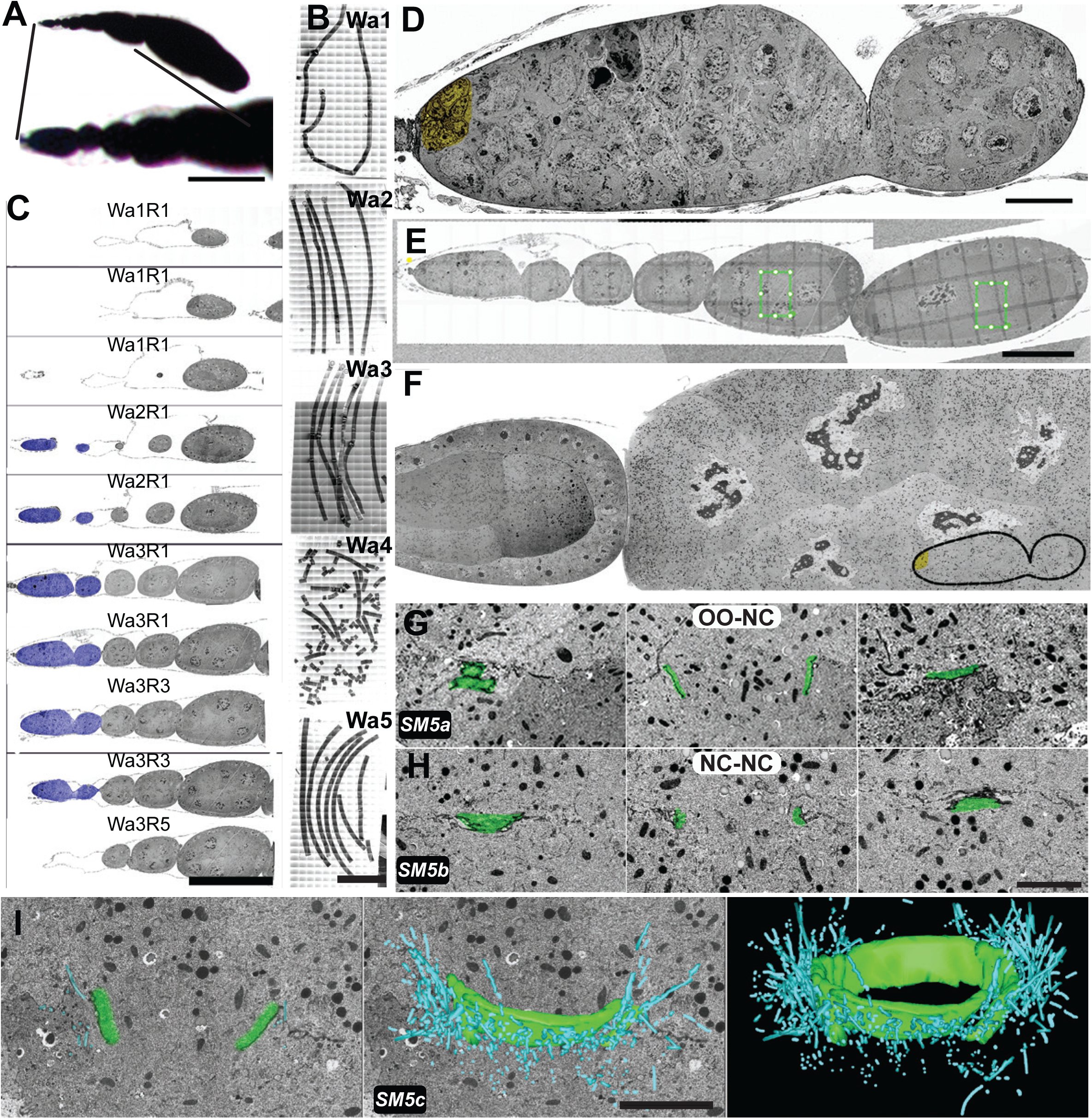
Array tomography and lateral screening can be used to locate and image ring canals within much larger, later stage egg chambers. (A) Image of an ovariole embedded after chemical preparation and darkened by the osmium treatment. Zoomed-in region represents the area that was sectioned in this experiment. Scale bar is 100 µm. (B) Ribbons of sections collected from the sample shown in (A) and attached to solid wafer supports. Serial sections are well-organized on wafers 1,2,3, and 5, but was disrupted on wafer 4; the order of the sections can be re-established during imaging or alignment. Scale bar is 1 mm. (C) Representative sections of the germarium (purple) were taken from different wafers and ribbons (indicated at the top of each image). Scalebar is 100 µm. (D) Single AT-SEM section through the germarium. The cap cells within the stem cell niche are pseudocolored yellow. Scale bar is 10 µm. (E) When the sample is properly oriented, a single AT-SEM section can capture an extensive region of the ovariole from the germarium through stage 7. Scale bar is 50 µm. (F) Single AT-SEM section showing a region of two older ECs within that ovariole. Bottom right corner shows the size of the germarium compared to this larger EC for scale (∼100 µm). (G,H) Three sections (first, middle, and last) of the same RC connect either a nurse cell to the oocyte (G) or two nurse cells to each other (H). Scale bar is 5 µm. (I) Rendering of the membranous interdigitations or “microvilli” (blue) and the actin-rich inner rim (green). Scale bar is 5 µm. SM refers to corresponding supplemental movies.

In addition to the actin-rich inner rim and the more electron-dense dark outer rim, a few studies have characterized the presence of highly convoluted membrane structures outside of the outer rim, which have been called microvilli (Loyer et al., 2015; Tilney et al., 1996). These interdigitations, which resemble structures termed junctions intercellular digitiform in the histology field, were previously only reported in later-stage ECs; they are proposed to anchor the RCs to the nurse cell membranes (Loyer et al., 2015; Tilney et al., 1996). Despite their essential role, these membrane structures have been challenging to study, and a detailed rendering of their three-dimensional structure has not been performed. Our AT-SEM images of RCs from mid-stage ECs were able to capture these extensive membrane interdigitations. Rendering of a targeted volume revealed that the interdigitations varied in length, and did not show an obvious coordination to their orientation (Fig. 5I, Supplemental Movies 5c). Future studies could determine whether the length or orientation of the interdigitations varies within an egg chamber based on location or lineage, and whether that changes during oogenesis.

Additional insight into the organization of the interdigitations was obtained from another AT-SEM sample in which the tissue structure was disrupted during the preparation process. In this sample, large gaps formed between nurse cells and between the nurse cells and the somatic follicle cell layer (Fig. 6A, arrows); these large gaps provided us with the opportunity to better visualize the interdigitations, which are typically tightly packed around the RCs. In this “imperfect” sample, we were able to capture multiple RCs in different orientations (Fig. 6B-D, Supplemental Movie 6), and we could clearly see membrane tubes within the tissue gaps. These structures were not as prevalent in regions of the sample that are further from the RCs (Fig. 6D; arrows), which suggests that their assembly or stabilization may be spatially controlled. Further, despite the gaps in the tissue, the actin-rich inner-rim and electron-dense outer-rim structure is still maintained (Fig. 6B-D), which is reminiscent of the RC detachment phenotype that has been reported in ECs with mutations in genes involved in membrane trafficking or adhesion (Coutelis and Ephrussi, 2007; Loyer et al., 2015; Murthy et al., 2005; Murthy and Schwarz, 2004; Oda et al., 1997; Peifer et al., 1993; Starble and Pokrywka, 2018; Tan et al., 2014; Vaccari et al., 2009; White et al., 1998). Additional work is required to determine the composition of this stable structure, especially the electron-dense outer rim, but it again highlights the value of using vEM to study the RC.

**Fig. 6:**
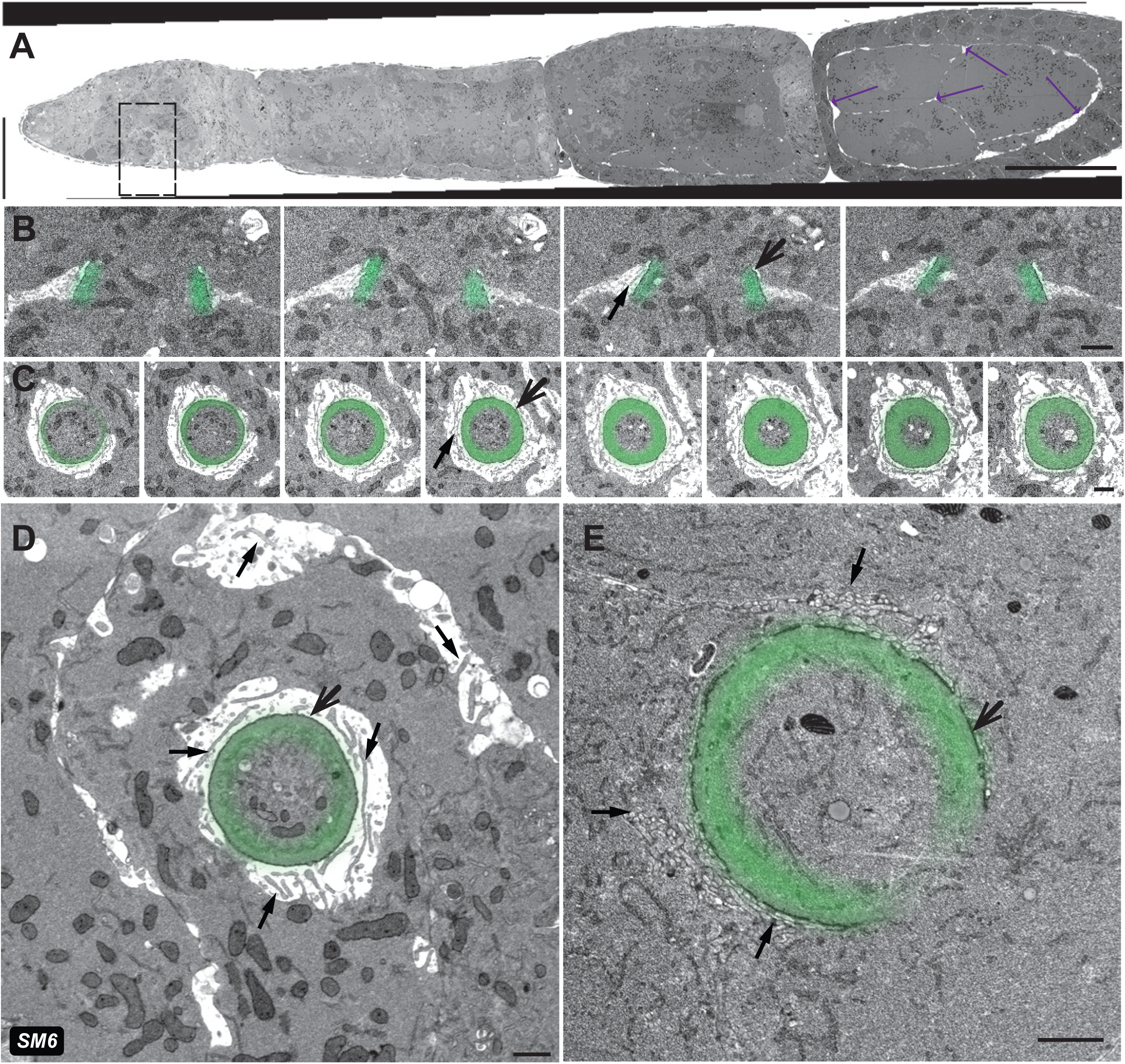
An imperfect sample revealed additional information about the tissue. (A) An AT-SEM image through the entire ovariole of an imperfectly preserved sample. Arrows point to gaps between cells in the tissue. Scale bar 50 µm. (B) A short z-series of images of two ring canals was obtained from this sample (5 nm lateral resolution). Membrane tubes, likely pieces of membrane interdigitations, can be seen in the gaps between cells. (C) A different sectioning orientation through a ring canal in the same sample. The long interdigitations are prominent around the RCs in all sections (black arrows). (D) Though the interdigitations are more concentrated close to the RCs, they are also present in the interface between the cells. (E) The dehydration artifact did not affect the ring canal from a different region sectioned in a similar orientation as the sample in panels C and D. The scale bars in panels B-E are 1 µm. SM refers to corresponding supplemental movie.

## DISCUSSION

The importance of vEM in cellular and developmental biology studies was highlighted recently (Collinson et al., 2023). Here, we have demonstrated that by using two complementary approaches, FIB-SEM and AT-SEM, we gained valuable insights into the three-dimensional ultrastructural changes that the germline RCs undergo during oogenesis. Our vEM data allowed us to monitor RC size, the thickness of the actin-rich inner rim, the structure of the nurse cell membranes, and the early appearance and orientation of extensive membrane interdigitations surrounding the RCs.

The extensive expansion of the germline RCs has been one of the reasons that this system has gained popularity in the study of intercellular bridges (Haglund et al., 2011; Robinson et al., 1994), but analysis of ultrastructural differences in RC size and thickness was lacking. Our data revealed significant differences in RC size and thickness based on developmental stage and lineage. Fluorescence-based analysis of RCs within the germarium suggested that the degree of contractile ring closure varies based on lineage, with the M1 contractile ring constricting to a diameter of 1.46 µm, while the M4 contractile ring constricts to a diameter of 0.79 µm. Further, there is evidence that RC growth may differ within the cluster; after the final mitotic division, the M1 canal is the first to initiate the growth phase (Koch and King, 1969; Ong and Tan, 2010). We have recently used fluorescence-based imaging to show lineage-based differences in ring canal scaling; the larger M1 RCs increase in size more slowly than the smaller RCs derived from subsequent mitotic divisions (Shaikh et al., 2024). These vEM data demonstrate that the thickness of the actin-rich inner rim differs between RCs within a cluster (Fig. 4, S4), which provides us with a potential mechanism to explain differences in the initiation and progression of RC growth, which could be further explored.

Our data also provided insights into the structure of the oocyte and nurse cell membranes. Previous TEM-based studies had described “microvilli” that surrounded the RCs in mid- to late-stage ECs (Fig.7A; Loyer et al., 2015; Tilney et al., 1996); these microvilli, which are likely stabilized by cadherin-based adhesions, were proposed to anchor the RCs within the nurse cell membranes (Loyer et al., 2015). Our data revealed that these extensive membrane interdigitations formed earlier than previously reported, at least by region 2b in the germarium.

**Fig. 7.**
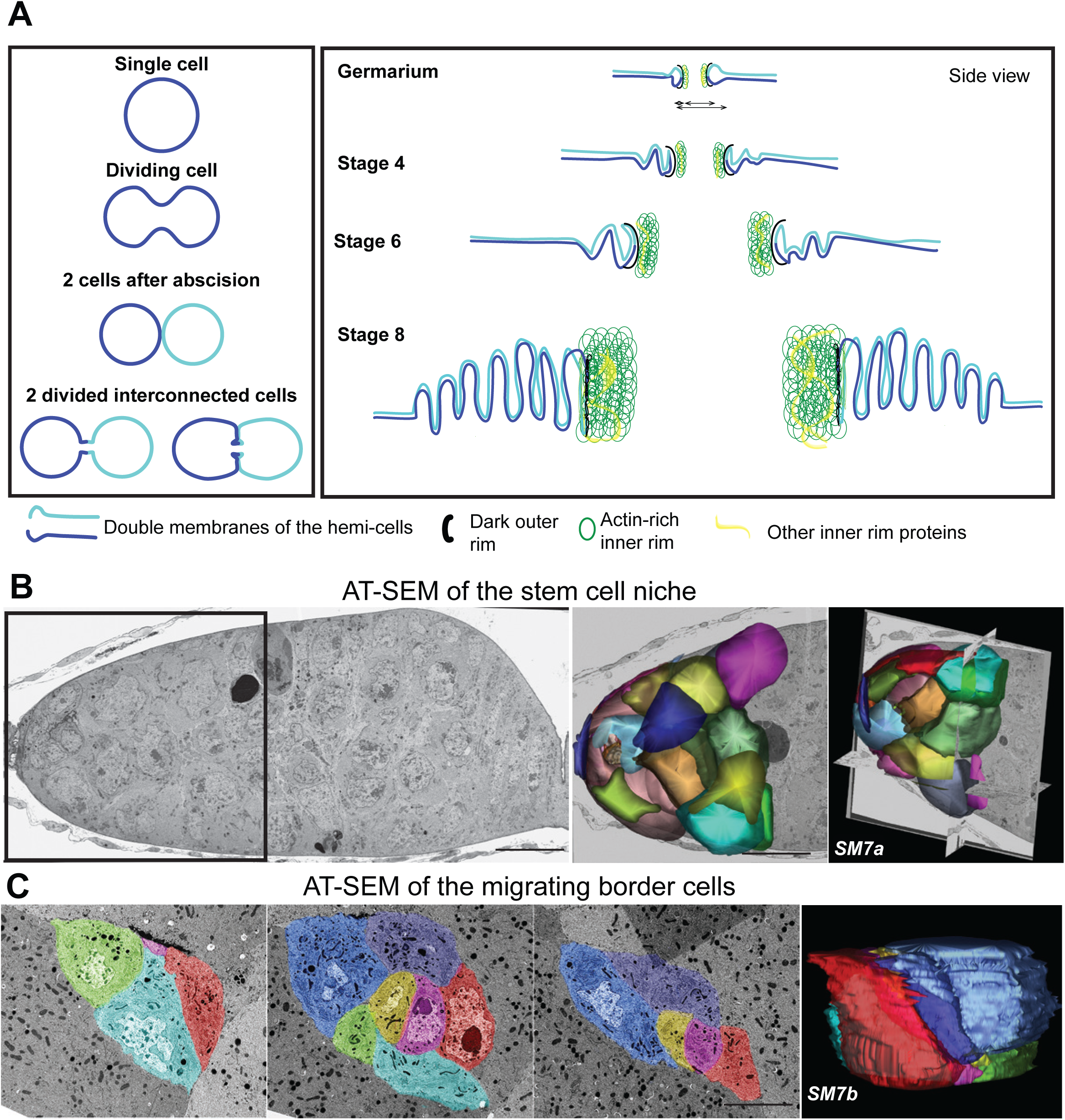
FIB-SEM and AT-SEM provide ultrastructural information about other cell types. (A) An illustration of the model of the incomplete cytokinesis of germline cells in the *Drosophila* egg chamber based on the ultrastructural insight provided by vEM imaging and published fluorescence data. Different cell membranes are represented in dark blue and turquoise. The division of ring canal parts is defined as the inner and outer rim. (B) AT-SEM image of the anterior end of the germarium that contains the stem cell niche. Serial sections were aligned and rendered to construct a 3D model of the cap cells. Scale bar 10 µm. (C) Serial sections of AT-SEM images were used to render a migrating border cell cluster model. Scalebar is 10 µm. SM refers to corresponding supplemental movies.

Although they could be seen along many parts of the nurse cell surface, they are more abundant near the RC (Fig. 3, 5, 6, 7A, S4). At later stages, these interdigitations became even more extensive and pronounced, and our volumetric AT-SEM data demonstrated that their length and orientation was not uniform (Fig. 5, 6, Supplemental Movie 6). The way that they extended from the RC into the oocyte-proximal nurse cell was reminiscent of the actin-based structures that were recently described (Lu et al., 2021), but a thorough characterization of these structures is required. For example, their formation and retraction could provide a way for the RCs to easily expand and contract to accommodate the transfer of different volumes of material or react to environmental changes. A recent study demonstrated that over-expression of the actin-binding protein, HtsRC, significantly increased RC size (Gerdes et al., 2020); therefore, future studies could determine whether HtsRC over-expression leads to more stable or more extensive interdigitations, which could provide a mechanism to increase RC growth or stabilize these larger RCs.

In addition to learning more about these intercellular bridges, our high-content datasets can provide ultrastructural insight into many other structures of interest, such as the fusome. Most analyses of the fusome have been done using a combination of fluorescence microscopy and 2D-TEM (Fig. 2A; (de Cuevas and Spradling, 1998; Deng and Lin, 1997; Grieder et al., 2000; Lighthouse et al., 2008; Lin et al., 1994; Lin and Spradling, 1995; Spradling et al., 1997), but our vEM dataset enabled us to generate a 3D model of this structure that provides a level of detail previously unavailable. Our rendering of the stage 2b clusters in the germarium supports a model for the gradual breakdown of the fusome (Fig. 3E), where the connected membrane network might break down first in the cluster’s periphery and then later in the central region. This fragmented pattern is consistent with prior live imaging studies that showed that by region 2b, the fusome was no longer continuous within the cluster (Snapp et al., 2004). Additional vEM analysis of the fusome could provide further insight into this important germline structure.

### AT-SEM can be used to capture small regions of the germarium or to locate “needle in a haystack” structures or cell types

AT-SEM can overcome the specimen size limitations that are encountered with FIB-SEM while providing a workflow to easily identify structures of interest within a large sample volume. In many cases, high-resolution imaging of an entire sample volume is unnecessary, and it would generate a large amount of data that must be stored and managed. With AT-SEM, a lower-resolution map of the sections is generated; from this map, a structure of interest can be quickly identified, and high-resolution imaging of a smaller, targeted volume can be performed. For example, our AT-SEM dataset contained sections of an entire ovariole from the stem cell niche and the mitotic region at the anterior to a stage 10 EC at the posterior. Because our FIB-SEM dataset did not capture the germline stem cell niche, we could easily image and render a small volume of this anterior region of the germarium to fill in this “gap” in our data. From this small volume, we generated a model in which we segmented the cap cells, which could be distinguished from other cells in the niche by their smaller size and electron-dense membranes (Spradling et al., 2008), as well as a dividing germline stem cell (Fig. 7B, Supplemental Movie 7a). Additional sections could easily be imaged to generate a 3D volume of the entire stem cell niche and the mitotic region of the germarium.

A known challenge in EM analysis is finding small, transient structures in a relatively large volume. Our AT-SEM dataset contained an EC at stage 9 of oogenesis, which is the stage in which the border cells are migrating. The border cells are a small group of somatic cells that delaminate from the follicular epithelium at the anterior end of the EC and collectively migrate through the germline cell cluster to the oocyte; these cells will ultimately give rise to the micropyle, which is the site of sperm entry (Rørth, 2002). The border cells have been used as a model for cancer cell metastasis (Stuelten et al., 2018), so learning more about the structure of these cells would improve our understanding of other types of invasive collective migration.

Although fluorescence microscopy of both fixed and live samples has revealed much about the dynamics of migration, because of their location deep within a large stage 9 EC, very little ultrastructural information exists. Lateral screening of the wafers allowed us to easily locate the migrating border cells, and we acquired high-resolution images from 60 sections and generated a simple model of the cluster (Fig. 7C, Supplemental Movie 7b). The ease with which these cells were located suggests that targeted AT-SEM analysis of multiple stage 9 ECs could provide a complete set of migration intermediates in controls and following different experimental manipulations, thereby improving our understanding of invasive cell migration.

### The potential of vEM data

vEM data can be generated using different experimental approaches, but each comes with its own set of strengths and limitations. By using a combination of FIB-SEM and AT-SEM, we have been able to capitalize on the strengths and overcome the limitations of each approach. FIB-SEM generates an easily aligned, high-resolution image stack, but because there is an upper limit on the size of the sample that can be imaged, this approach could not be used to monitor ultrastructural changes in larger samples. With AT-SEM, images of much larger sample surfaces can be acquired hierarchically: lower-resolution images can be collected initially to provide a map for screening, and then higher-resolution images can be collected of the relevant structures (Fig. 5). Once a region of interest is identified in one section, it is straightforward to image sections above and below to capture a small 3D volume of that part of the sample. With larger samples and higher resolution imaging, it is difficult to accurately estimate the dimensions of the acquired data: the germarium, which is larger than the volume attainable by FIB-SEM, constitutes only a small fraction of the stage 9 EC (Fig. 5F, trace to scale). Therefore, attempting to collect the entire volume of such a large structure at high resolution is impractical. However, because the sections containing the relevant structures of interest can be easily located, volume data that covers just the relevant sections can be acquired; this significantly reduces acquisition time and eases the burden on computational resources. If additional structures become of interest later, the low-resolution map can be used to easily locate and acquire images of a new region of interest.

By using EM, the researcher can monitor the location and abundance of the entire morphome of the cell or tissue of interest. Recently described as the “silent revolution” in electron microscopy, vEM data provides ultrastructural insight that would be impossible to glean from any single-section EM approach (Collinson et al., 2023). As we have demonstrated, this approach provides not only valuable insight into the structure of interest, but also supplies ultrastructural information about features that were either not expected (such as the membrane lining the RC lumen) or not the primary focus (the fusome, migrating border cells, and stem cell niche). Such high-content datasets can foster new collaborations among researchers studying diverse questions within the same system and promote scientific equity by providing access to datasets that might be out of reach for many researchers to generate themselves due to limited funding or access to the necessary equipment or expertise.

## ABBREVIATIONS

2D: Two Dimensional
3D: Three Dimensional
AT: Array Tomography
EC: Egg Chamber
EM: Electron Microscopy
FIB: Focused Ion Beam
HPF: High-pressure Freezing
RC: Ring Canal
ROI: Region of interest
SBF: Scanning Block Face
SEM: Scanning Electron Microscopy
TEM: Transmission Electron Microscopy
vEM: Volume Electron Microscopy
X, Y, Z,: special axis designation

## FIGURE LEGENDS

**Figure S1.**
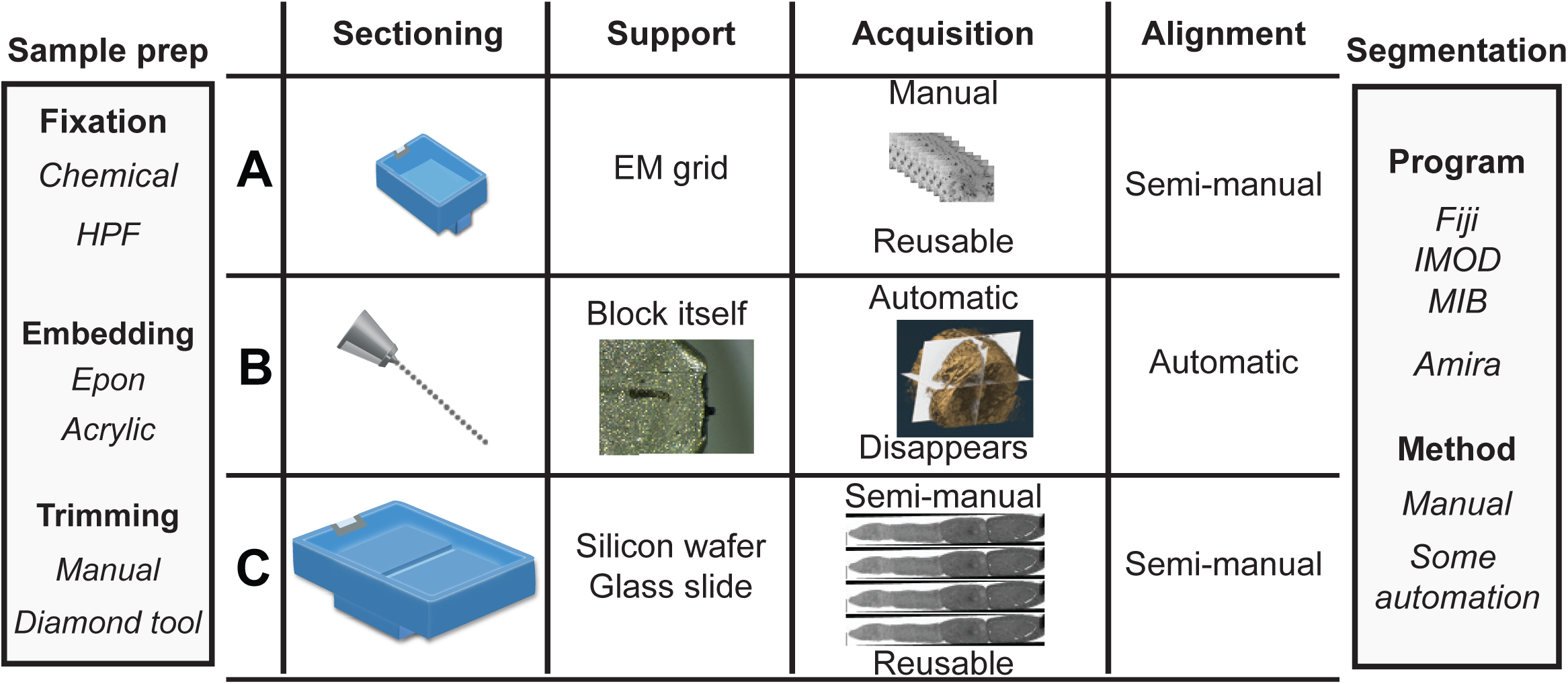
Sample preparation, acquisition, alignment, and analysis for different types of vEM. All described techniques are performed at room temperature with adaptations in preparation techniques based on the type of the sample. (A) For TEM, the samples are sectioned on an ultramicrotome with a diamond knife and transferred to support grids (round, 3mm diameter, thin metal foil). The maximum area of the sample is limited by these dimensions. The thickness of the sections varies between 50 and 300 nm and is defined by the penetration capacity of the electron beam. The resulting images provide a very high resolution (∼ 0.2 nm), but higher magnification images result in a smaller field of view. (B) For FIB-SEM, minimal manipulation is required before introducing the sample into the SEM sample chamber. The block surface is milled directly inside the SEM chamber and the images are collected from the newly exposed surface. The destructive nature of the acquisition prevents modification or reacquisition of the images once acquired. FIB removes 5-50 nm of the surface before capturing a SEM image, with the cycle continuing to create a nearly perfect 3D stack through a relatively small sample volume. The stack of images is easy to align, and isotropic resolution can be achieved. (C) For AT-SEM, blocks are sectioned using a diamond knife, creating a sequence of sections (arrays), which is transferred to a large support, such as a wafer or a coated glass coverslip. The physical sectioning by the diamond knife permits the processing of a relatively large area (> 1 mm^2^) and a nearly unlimited z depth. The primary strength of the technique is the ability to use lateral screening of the arrays to efficiently locate the ROI, which can then be selectively imaged at higher resolution. With tiled acquisition there is no real upper limit to the area that can be acquired; further, sections are stable and can be repeatedly imaged at a range of resolutions. z resolution is limited by physical sectioning to ∼50 nm. The alignment and segmentation steps of the workflow are frequently performed with the same program. The alignment can be performed automatically with the help of the program’s algorithms or manually by adjusting the consecutive images individually. The segmentation depends on the scope of the desired model and the abundance and complexity of the structures; it can be done manually or with automation.

**Supplemental Movie 1a**. Movie of entire FIB-SEM image stack through the germline clusters within the germarium, which are rendered in Fig. 3. https://doi.org/10.6084/m9.figshare.28430189

**Supplemental Movie 1b**. Movie of entire FIB-SEM image stack through part of the stage 4 egg chamber. https://doi.org/10.6084/m9.figshare.28430192

**Supplemental Movie 2**. Movie of a FIB-SEM serial section alignment through the ring canal presented in Fig. 2B, C. https://doi.org/10.6084/m9.figshare.28430066

**Supplemental Movie 3a.** A series of aligned sequential EM sections was obtained using FIB-SEM, focusing on the germline region. Different clusters were manually segmented with separate colors for each cluster. Ring canals for each cluster were generated using the isosurface feature of IMOD. https://doi.org/10.6084/m9.figshare.28430096

**Supplemental Movie 3b.** A series of sequential EM sections were obtained using FIB-SEM, focusing on the stage 2b purple cluster. The segmentation highlights the complexity of the fusome in the germline (white) and its interaction with the ring canals connecting different germline cells within the cluster. https://doi.org/10.6084/m9.figshare.28430117

**Supplemental Movie 4.** Aligned sequence of the RCs from the FIB-SEM dataset of the stage 4 EC shown in Fig. 4A. The inner rim of the ring canals is pseudocolored (green). https://doi.org/10.6084/m9.figshare.28430213

**Fig. S4:**
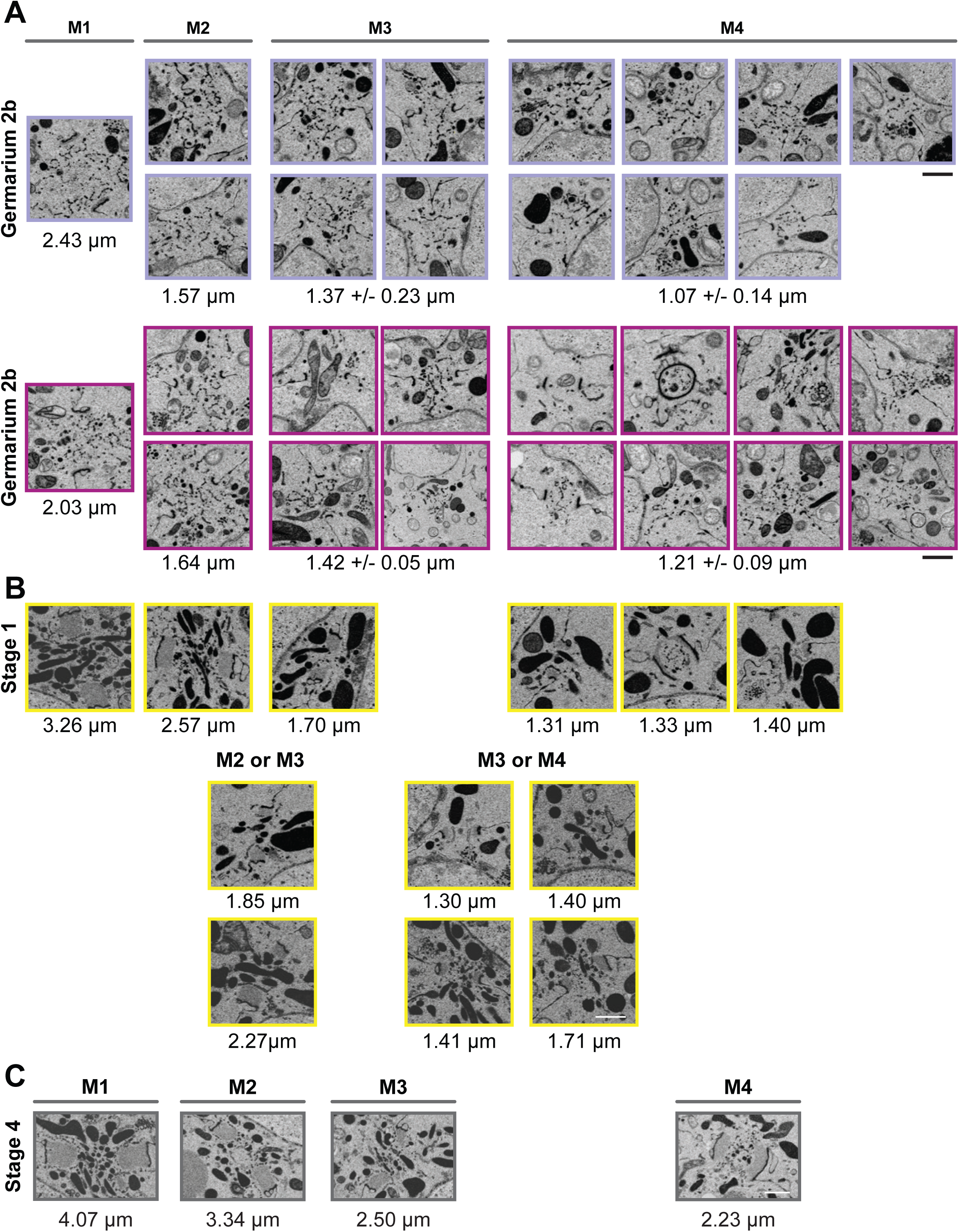
Single-plane FIB-SEM images show the differences in ring canal size, thickness, and orientation within each cluster. Images were cropped to highlight the germline RCs within each cluster. (A) The two clusters from region 2b (purple and pink) were sufficiently covered to determine mitotic division of origin for all RCs. (B) For the stage 1 cluster (yellow), division of origin could be definitively assigned for about half of the RCs, and for the other half, origin could be narrowed down to one of two types. (C) The older, stage 4 EC (grey) was not completely captured, so only the four posterior RCs directly connecting to the oocyte could be definitively characterized. Scale bars are 1 µm.

**Supplemental Material 5.** A composite of tiled images acquired with 5 nm resolution. Uncompressed image represents 1101×266 cm. https://doi.org/10.6084/m9.figshare.28430216

**Supplemental Movie 5a.** Aligned serial sequence of the OO-NC RC from the AT acquisition shown in Fig. 5G. https://doi.org/10.6084/m9.figshare.28430138

**Supplemental Movie 5b.** Aligned serial sequence of the NC-NC RC from the AT acquisition shown in Fig. 5H. https://doi.org/10.6084/m9.figshare.28430147

**Supplemental Movie 5c.** Aligned serial sequence of the OO-NC RC from the AT acquisition shown in Fig. 5G. The movie combines the EM sequence and the subsequent segmentation of the outline of the ring canal (green) and the portions of the interdigitations (light blue). https://doi.org/10.6084/m9.figshare.28430258

**Supplemental Movie 6:** Serial sections through the RC from the imperfectly prepared sample shown in Fig. 6. https://doi.org/10.6084/m9.figshare.28430162

**Supplemental Movie 7a:** Aligned serial sections of AT-SEM images of the anterior end of the germarium containing the germline stem cell niche. The movie combines the EM sequence and the subsequent segmentation. The image stack shows what appears to be a dividing germline stem cell; the two daughter cells are pseudo-colored. https://doi.org/10.6084/m9.figshare.28430165

**Supplemental Movie 7b:** Aligned serial sections of AT-SEM images. The movie combines the AT-SEM sequence and the subsequent segmentation of the border cell cluster. https://doi.org/10.6084/m9.figshare.28430177

## Acknowledgements

We would like to thank Rebecca Green, Fani Papagiannouli, David Hall, Fanny Langlet, and Gareth Griffiths for the critical reading of the manuscript and helpful suggestions. We thank Helmut Gnaegi of Diatome for the constant support in AT diamond knife improvement. This work was supported by the National Institutes of Health (NIH R15HD084243 to LL). The purchase of the confocal microscope used in this study was supported by NSF-MRI Award #2116348. Stocks obtained from the Bloomington *Drosophila* Stock Center (NIH P40OD018537) were used in this study. The following antibody was obtained from the Developmental Studies Hybridoma Bank, created by the NICHD of the NIH and maintained at The University of Iowa, Department of Biology, Iowa City, IA 52242: hts RC antibody developed by L. Cooley, hts (1B1) developed by H.D. Lipshitz. The IMOD software is supported by NIH/NIGMS grant GM125074 to David Mastronarde.

